# A monochromatically excitable green-red dual-fluorophore fusion incorporating a new large Stokes shift fluorescent protein

**DOI:** 10.1101/2023.07.16.549156

**Authors:** J. Obinna Ejike, Mayuri Sadoine, Yi Shen, Yuuma Ishikawa, Erdem Sunal, Sebastian Hänsch, Anna B. Hamacher, Wolf B. Frommer, Michael M. Wudick, Robert E. Campbell, Thomas J. Kleist

## Abstract

Genetically encoded sensors enable quantitative imaging of analytes in live cells. State-of-the-art sensors are commonly constructed by combining ligand-binding domains with one or more sensitized fluorescent protein (FP) domains. Sensors based on a single FP are susceptible to artifacts caused by differing expression levels or sensor distribution *in vivo*. Hence, our lab developed dual-FP Matryoshka technology introduced by a single cassette that contains a stable large Stokes shift (LSS) reference FP nested within a reporter FP (cpEGFP), allowing simple construction of intensiometric sensors with the capacity for ratiometric quantification. The first-generation Green-Orange (GO) Matryoshka cassette established proof of concept but required custom optical setups to maximize achievable dynamic range. Here, we present a genetically encoded calcium sensor that employs optimized second-generation Green-Apple (GA) Matryoshka technology that incorporates a newly designed red LSSmApple fluorophore. LSSmApple provides improved excitation spectrum overlap with cpEGFP, allowing for monochromatic co-excitation with blue light. The exceptionally large Stokes shift of LSSmApple results in improved emission spectrum separation from cpEGFP, which minimizes fluorophore bleed-through and facilitates imaging using standard dichroics and red fluorescent protein (RFP) emission filters. We developed an image analysis pipeline for yeast (*Saccharomyces cerevisiae*) timelapse imaging that utilizes LSSmApple to segment and track cells for high-throughput quantitative analysis. In summary, we engineered a new fluorescent protein, constructed a genetically encoded calcium indicator (GA-MatryoshCaMP6s), and performed calcium imaging in yeast as a demonstration.

## Introduction

At the subcellular level, biological processes occur on a scale that is not visible to the human eye. In the late 20th century, the identification and application of the *Aequorea victoria* jellyfish green fluorescent protein (avGFP) revolutionized cell biology research by enabling *in vivo* observation of cellular processes previously unseen^1–3^. Since then, researchers have created a diverse range of fluorescent proteins (FPs) with enhanced features such as superior quantum yield, photostability, and distinctive emission spectra^4,5^. One important spectral property of FPs is their Stokes shift, which is described as the distance in the wavelength between their absorption and emission maxima^6^. FPs with a Stokes shift ≥ 100 nanometers (nm) are typically classified as large Stokes shift (LSS) FPs. LSS FPs can be advantageous in multiplex imaging, as they allow simultaneous monochromatic excitation but provide spectral separation from commonly used non-LSS FPs such as enhanced GFP (EGFP). For many applications, ideal LSS red FPs (LSS RFPs) would have similar excitation spectra to EGFP. Excitation at a single wavelength increases the maximum achievable imaging speed and reduces phototoxicity and photobleaching. Further, LSS FPs reduce the setup costs for multiplex imaging in two-photon microscopy, which requires expensive (e.g., femtosecond titanium-sapphire) lasers for excitation^7–9^. Genetically encoded sensors have been developed using FPs, which can spatiotemporally report on analyte concentrations *in vivo*^10–13^. These sensors can be categorized as single- or dual-FP sensors. Single-FP sensors can exploit either their inherent sensitivity to specific stimuli or can be fused with a recognition element that responds to a specific process or analyte. Circular permutation (cp), in which the N- and C-termini of a FP are interchanged and reconnected by a small linker, can help to sensitize the FP to conformational changes in a fused recognition element^14,15^. Dual-FP sensors often utilize Förster resonance energy transfer (FRET). FRET is a non-radiative energy transfer in which a donor chromophore transfers energy to an acceptor chromophore. FRET requires overlap of the donor emission and the acceptor excitation spectra. Further, FRET efficiency is highly sensitive to the proximity and relative orientation of donor and acceptor chromophores. FRET sensors commonly consist of a recognition domain sandwiched between two FRET compatible FPs. Conformational changes in the recognition domain result in changes in the proximity and orientation of the FRET pair and concomitant changes in FRET efficiency^10,15^.

Single-FP intensiometric sensors exploit conformational changes that affect the extinction coefficient and/or quantum yield of the chromophore, thereby altering fluorescence intensity. Empirically, state-of-the-art intensiometric single-FP sensors have been found to outperform FRET-based sensors in terms of dynamic range and signal-to-noise ratio (SNR), however intensiometric sensors are susceptible to artifacts derived from variable expression levels, bleaching, or sample movement. Dual-FP FRET sensors, on the other hand, are less prone to these artifacts as they allow a ratiometric readout. Due to the size of the FP barrels, which restricts proximity of the chromophores, dual-FP FRET sensors usually cannot reach maximal theoretical FRET efficiencies between the two chromophores, which results in lower realized dynamic ranges than single FP sensors^16,17^. Previously, our group developed a new sensor technology, called Matryoshka, which nests a reference LSS FP within a cpEGFP for simple conversion of cpEGFP-based intensiometric sensors into dual-FP ratiometric sensors^16^. In this study, to create a ratiometric cassette for high SNR imaging, we engineered a novel red LSS-FP, with a high excitation spectral overlap with cpEGFP, resulting in an improved Green-Apple (GA) Matryoshka cassette. The optimized GA fluorescent Matryoshka cassette was used to generate a new calcium sensor, GA-MatryoshCaMP6s, that enables facile ratiometric imaging with high dynamic range and SNR. GA-MatryoshCaMP6s was applied for calcium imaging of the alpha-mating factor (α-MF) response in budding yeast (*Saccharomyces cerevisiae*).

## Results

### Engineering and directed evolution of a new large Stokes shift red FP

To engineer a bright LSS RFP with efficient excitation at ∼488 nm illumination, we used a bright and photostable RFP, monomeric Apple (mApple), as a template and created a library of variants carrying all possible proteinogenic amino acids at residues 161 and 163 (numbered according to positions in the parental FP DsRed). It was previously reported that an acidic residue at positions 161 or 163 in RFP could potentially serve as a proton acceptor for the protonated chromophore. Establishing this excited state proton transfer (ESPT) pathway around the protein chromophore allows LSS emission to occur^9,18^. *E. coli* colonies expressing the mutant library were visually screened under blue light for yellow fluorescent color, with the rationale that the strong blue absorbance of the protonated chromophore would lead to yellow-hued colonies. Positive colonies were picked and cultured for extraction of protein variants, and fluorescence spectra were measured. Some variants exhibited prototypical LSS fluorescence with distinctive blue excitation and red emission. Sequencing of candidate mutants revealed that one of the residues in positions 161 and 163 was mutated to either glutamic acid (E) or aspartic acid (D), including I161E/K163G, I161G/K163D, I161C/K163E, and I161S/K163E. With the successful establishment of LSS fluorescence, we sought to further improve the brightness of the mutant LSS RFP prototypes by combining semi-rational design and random mutagenesis. Saturation mutagenesis was performed at positions 143, 146, 161, and 163, which are predicted to be in close proximity to the protein chromophore. We then performed several rounds of random mutagenesis and brightness screening, leading to the identification of a final variant that was designated LSSmApple (Figure 1A).

**Figure 1.**
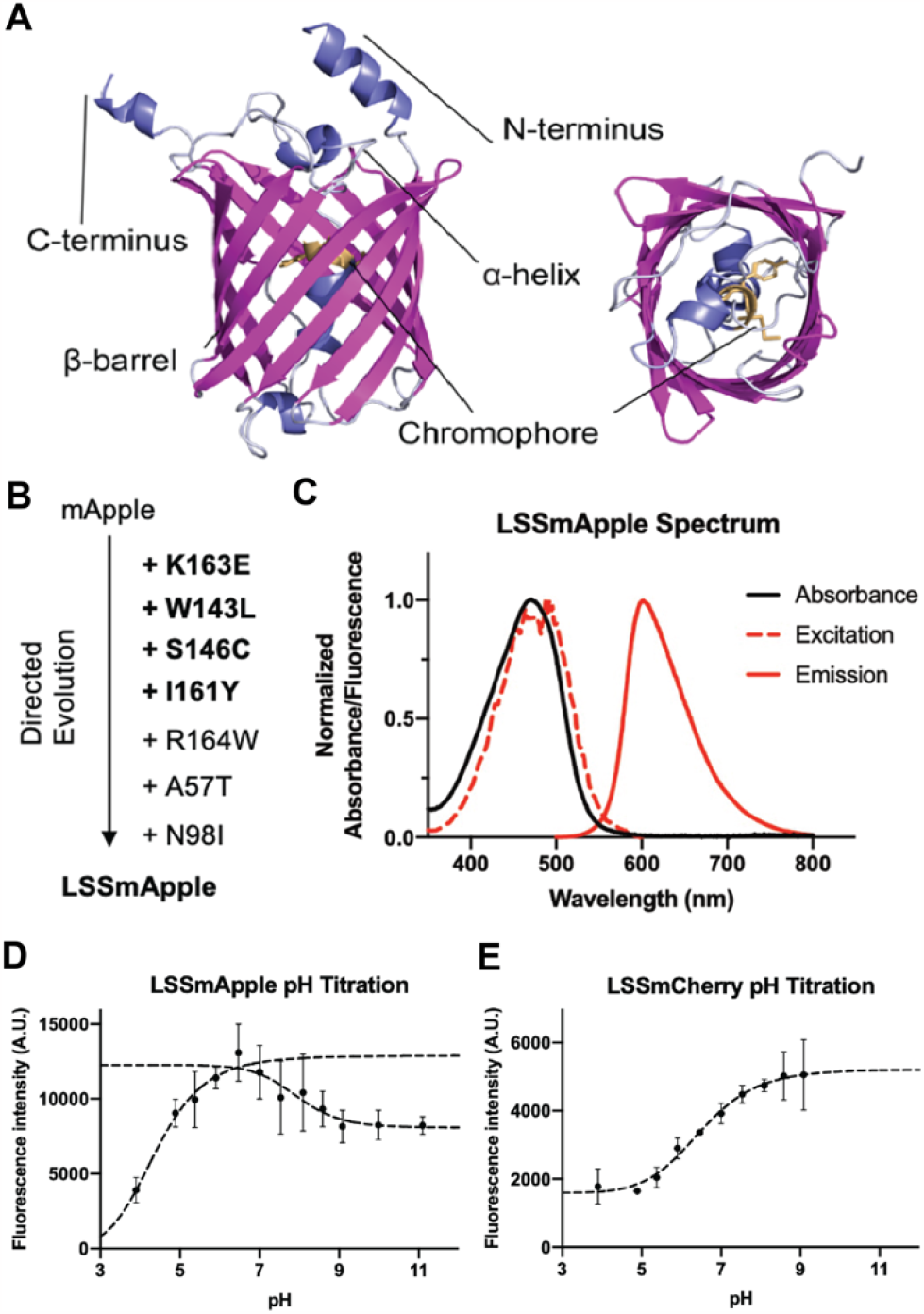
Engineering and characterization of LSSmApple. **(A)** Model of the LSSmApple FP. The panel on the left side shows the β-barrel of LSSmApple, while the right panel shows the chromophore. For better visualization of the chromophore, N- and C-terminal α-helices were hidden in right panel. See confidence of model in Figure S1 **(B)** Evolution chart of amino acid substitutions resulting in LSSmApple. Mutations from site-directed saturation mutagenesis are in bold. Mutations are listed in the chronological order. **(C)** Absorbance (solid black line), excitation (red dotted line), and emission (red solid line) spectra of LSSmApple. **(D)** *In vitro* pH titration curves for LSSmApple. The pH ranges 4-7 and 7-11 were fitted separately. **(E)** *In vitro* pH titration curve for LSSmCherry, as a comparison. Errors bars indicate standard deviation (n=3).

LSSmApple has seven mutations compared to its ancestral mApple variant: A57T / N98I / W143L / S146C / I161Y / K163E / R164W (Figure 1B). With an excitation peak at 465 nm and emission peak at 600 nm, LSSmApple has a large Stokes shift of 135 nm (Figure 1C). LSSmApple has an extinction coefficient of 45,000 M^-1^cm^-1^, and a quantum yield of 0.31, which results in ∼40% greater brightness compared to our previously published LSSmCherry1^9^. Titrations of pH for LSSmApple indicated its LSS fluorescence is stable over the physiological pH range (pH 5 – pH 8) with a pK_a_ at 4.3, whereas LSSmCherry1 is relatively pH sensitive, with a pK_a_ at 6.2 (Table 1, Figure 1D-E).

**Table 1.** Biophysical properties of the LSS FPs tested as stable reference FPs in Matryoshka cassettes.

|  | Extinction coefficient<br>(EC, M<sup>-1</sup>cm<sup>-1</sup>) | Quantum yield<br>(QY) | Brightness (EC x<br>QY, M<sup>-1</sup> cm<sup>-1</sup>) | Excitation max.<br>(nm) | Emission max.<br>(nm) | pK<sub>a</sub> | Maturation t<sub>1/2</sub> *<br>(min) |
| --- | --- | --- | --- | --- | --- | --- | --- |
| LSSmOrange<sup>8</sup> | 52000<sup>8</sup> | 0.45<sup>8</sup> | 23.4<sup>8</sup> | 437<sup>8</sup> | 572<sup>8</sup> | 5.7<sup>8</sup> | 138<sup>8</sup> |
| LSSmCherry1<sup>9</sup> | 35000<sup>9</sup> | 0.29<sup>9</sup> | 10<sup>9</sup> | 450<sup>9</sup> | 610<sup>9</sup> | 6.2<sup>9</sup> | 90<sup>9</sup> |
| LSSmApple | 45000 | 0.31 | 14 | 465/490 | 600 | 4.3/7.9 | 110 |
| * |  | At |  | 37 |  | °C |  |

### Design of Green-Apple Matryoshka cassette

To investigate whether LSSmApple could function as a suitable reference for a Matryoshka cassette, we compared its spectral properties to the previously published LSSmCherry1^9^ and LSSmOrange^19^, which is the reference fluorophore in GO-Matryoshka^16^. LSSmApple and LSSmChery showed greater emission spectra separation from cpEGFP compared to LSSmOrange (Figure 2A-D). Each of the red FPs showed shorter maturation times than LSSmOrange (Table 1). Of the three LSS FPs, LSSmApple has the greatest excitation spectra overlap with cpEGFP, showing two peaks at 465 and 490 nm. LSSmApple has a greater extinction coefficient and quantum yield, thus LSSmApple is brighter than LSSmCherry1 and shows better pH stability (pK_a_=4.3/7.9) (Table 1). Hence, LSSmApple was chosen as a reference for a monochromatically excitable Green-Apple Matryoshka cassette (Figure 2E).

**Figure 2.**
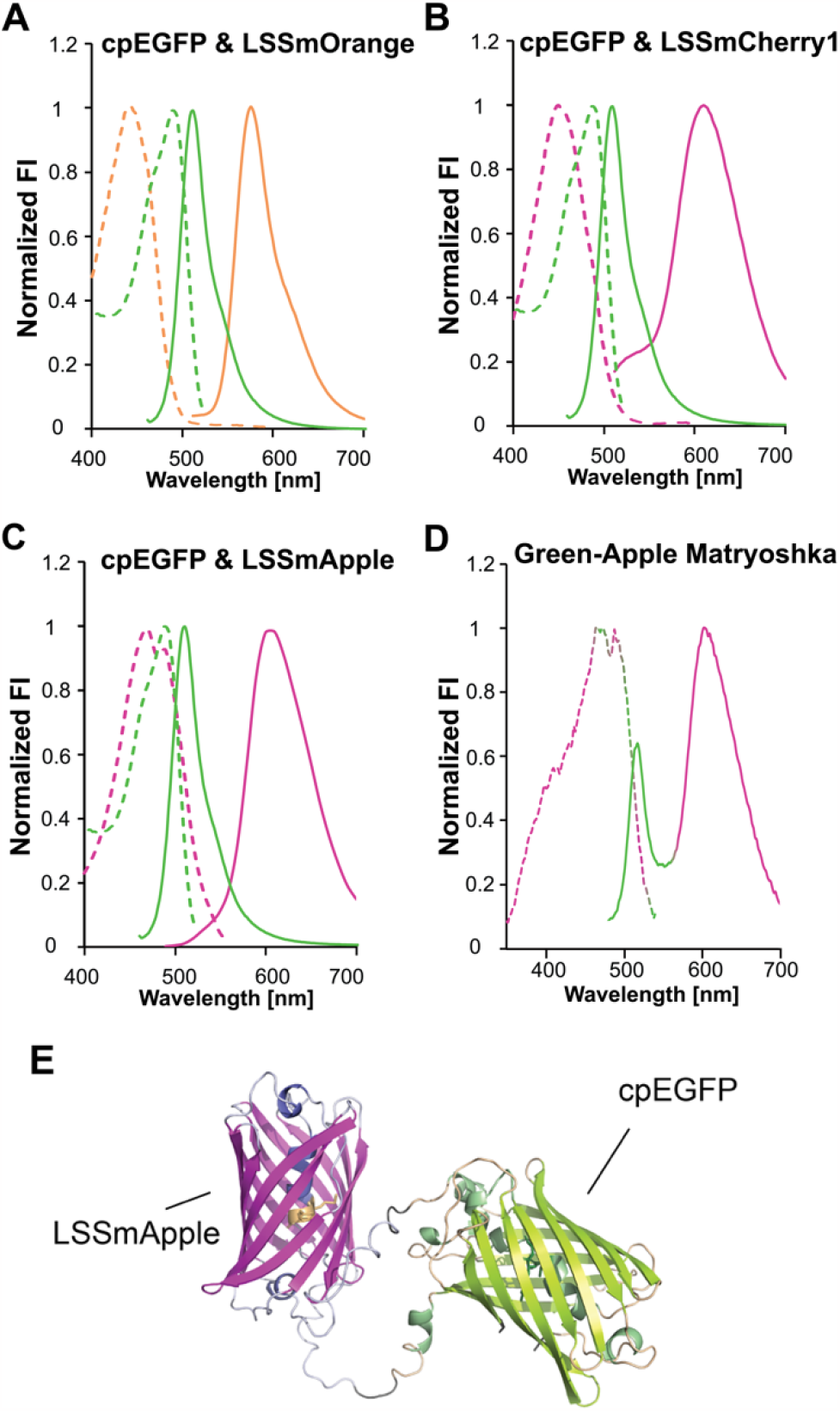
Comparison of spectral properties of FPs for the generation of Green-Red Matryoshka cassette. (A) Excitation (dotted line), and emission (solid lines) spectra of LSSmOrange (B) LSSmCherry1 (C) LSSmApple relative to cpEGFP. (D) Spectral characteristics of the Green-Apple Matryoshka cassette (E) AlphaFold2 model of the Green-Apple Matryoshka cassette. See confidence of model in Figure S1. FI: Fluorescence intensity.

### Creation and characterization of GA-MatryoshCaMP6s

LSSmApple was chosen as reference for the second-generation GA-Matryoshka cassette due to its greater spectral overlap with cpEGFP in comparison to LSSmCherry1 and LSSmOrange. As proof of concept, we turned to the scaffold of the genetically encoded calcium indicator (GECI) GCaMP6s^20^, which is an ultrasensitive calcium sensor consisting of a cpEGFP sandwiched between a myosin light chain kinase-derived calmodulin binding peptide (RS20) and a calmodulin (CaM) (Figure 3A-C). Previously, we had used this sensor scaffold with LSSmOrange to create GO-MatryoshCaMP6s^16^. *In vitro* characterization of purified GA-MatryoshCaMP6s revealed two emission maxima at λ_em_ ∼515 nm and λ_em_ ∼600 nm when excited at λ_ex_ 453 nm (Figure 3D). Titration of increasing concentrations of free calcium demonstrated a positive correlation with cpEGFP fluorescence (Figure 3D). LSSmApple fluorescence (∼600 nm) increased slightly at higher concentrations of calcium, which might be a result of fluorescence bleed-through or FRET. We calculated a dissociation constant (K_d_) of 101 ± 3 nM (mean ± s.e.m., n=3) for GA-MatryoshCaMP6s (Figure 3 D), which is similar to reported values for GCaMP6s^16^, and a ratiometric dynamic range (ΔR/R_0_) of 27.60 ± 0.82 (mean ± s.e.m.) for GA-MatryoshCaMP6s. Thus, in vitro characterization of GA-MatryoshCaMP6 demonstrated that the GA-Matryoshka cassette can be used to easily convert intensiometric cpEGFP-based sensors into dual-fluorophore sensors capable of ratiometric measurements with high dynamic range using monochromatic illumination.

**Figure 3.**
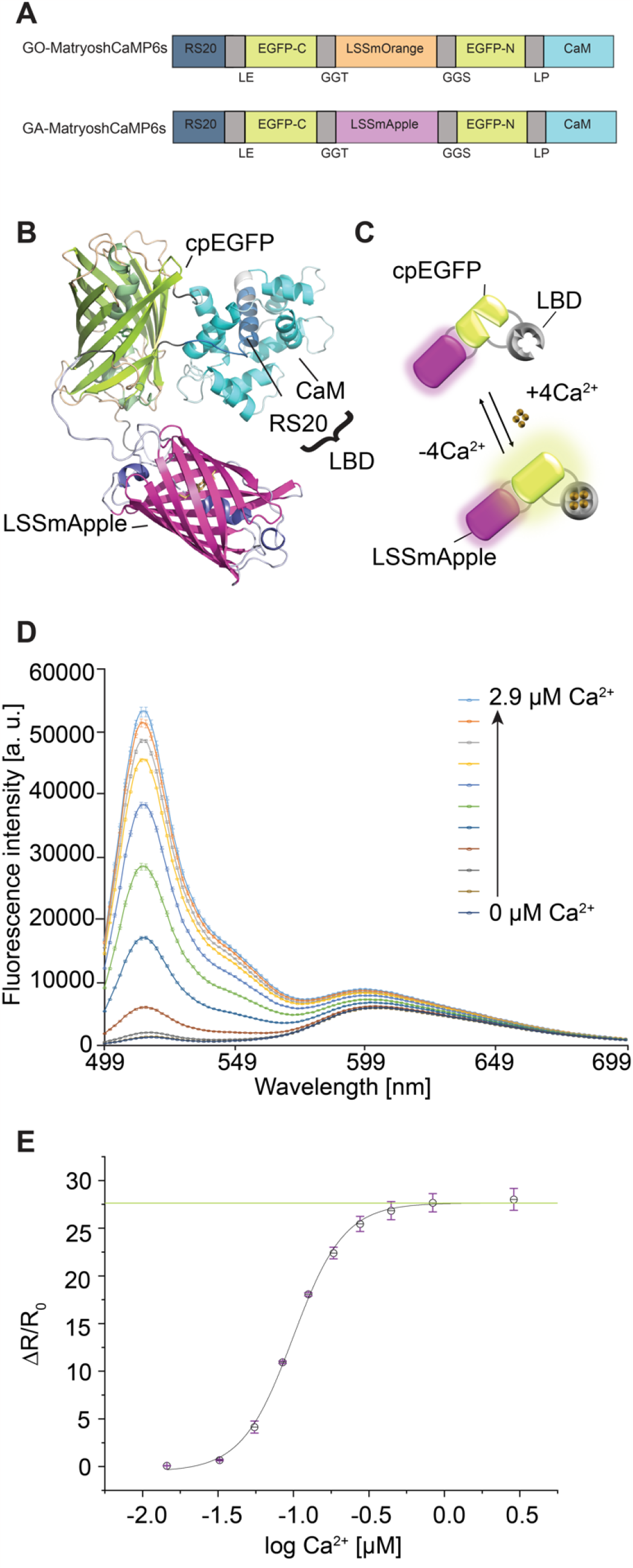
Green-Apple MatryoshCaMP6s design and *in vitro* characterization. **(A)** Schematic representation of GO-MatryoshCaMP6s and GA-MatryoshCaMP6s. **(B)** AlphaFold2 structure prediction of the GA-MatryoshCaMP6s. The sensor is a fusion of four polypeptides: the cpEGFP (green), a bipartite ligand binding domain (LBD) from CaM-RS20 (dark blue and light-blue), and the reference LSSmApple (magenta). Linkers shown in grey. See confidence of model in Figure S1. **(C)** Cartoon depicting the mechanism of GA-MatryoshCaMP6s. **(D)** Steady-state emission spectra of GA-MatryoshCaMP6s at 11 different calcium concentrations. Mean of n=3 technical replicates plotted. Error bars indicate s.e.m. The two other biological replicates showed comparable results. **(E)** Titration curve of GA-MatryoshCaMP6s from n=3 biological replicates in two independent titrations. Error bars indicate s.e.m.; green line indicates dynamic range of GA-MatryoshCaMP6s (ΔR/R_0_). Sigmoid fit done *via* Boltzmann function.

### Calcium imaging in yeast with GA-MatryoshCaMP6s

In budding yeast (*Saccharomyces cerevisiae*), calcium signaling coordinates responses to osmotic stress, glucose, and mating pheromones^21,22^. We used α-mating factor (α-MF) to elicit calcium spiking in yeast cells with mating type a (*MATa*) expressing GA-MatryoshCaMP6s and monitored cytosolic spiking calcium dynamics using trapped-cell microfluidics and spinning disk confocal microscopy^23^. Yeast were pre-grown in microfluidic chambers overnight, which resulted in enabling us to visualize hundreds of cells in a single field of view (FOV). We imaged yeast for approximately 10 min while perfusing growth medium before switching to growth medium supplemented with 50 μM α-MF for the remainder of the 140-minute experiment. A far-red fluorescent dye was used as a tracer to mark and time the arrival of the medium containing α-MF. We created a software pipeline that used the LSSmApple channel to segment individual cells, track them, and extract fluorescence intensities and spatiotemporal parameters. Stable fluorescence from LSSmApple facilitated tracking and segmenting the moving cells during the experiment. Tracking and segmentation would be difficult using cpEGFP fluorescence, which is dim at resting state and changes several-fold in intensity during calcium elevations. Yeast cells were initially perfused with growth medium. After 10 min, medium with 50 μM α-MF was perfused for 120 min. Cytosolic calcium spikes were observed to occur in an asynchronous manner across the population of cells, and spiking frequency increased after treatment with α-MF (Figure 4A and 4B, Supplemental Figures S2 and S3). After ∼50 min (∼40 min lag time after arrival of α-MF) a noticeable increase in spiking frequency could be observed across the full FOV (Figure 4C, Supplemental Figure S2). In the two hours of treatment, single cells spiked approximately 5 times on average (n=409) (Figure 4D). Calcium spikes were asymmetric, with short rise times and slower returns to the baseline. Rise times were quantified for all spikes (n=1898) and were at average 0.54±0.04 min, while spike fall times were noticeably longer, with an average of 1.14±0.02 min (Figure 4E). The average maximum amplitude of all spikes was calculated as a ratio change (cpEGFP / LSSmApple) of 3.18±0.03 (mean ± s. e. m.) (Figure 4F). Collectively, the results demonstrate the utility of GA-MatryoshCaMP6s, including the LSSmApple reference which enables segmentation and tracking, for *in vivo* calcium imaging.

**Figure 4.**
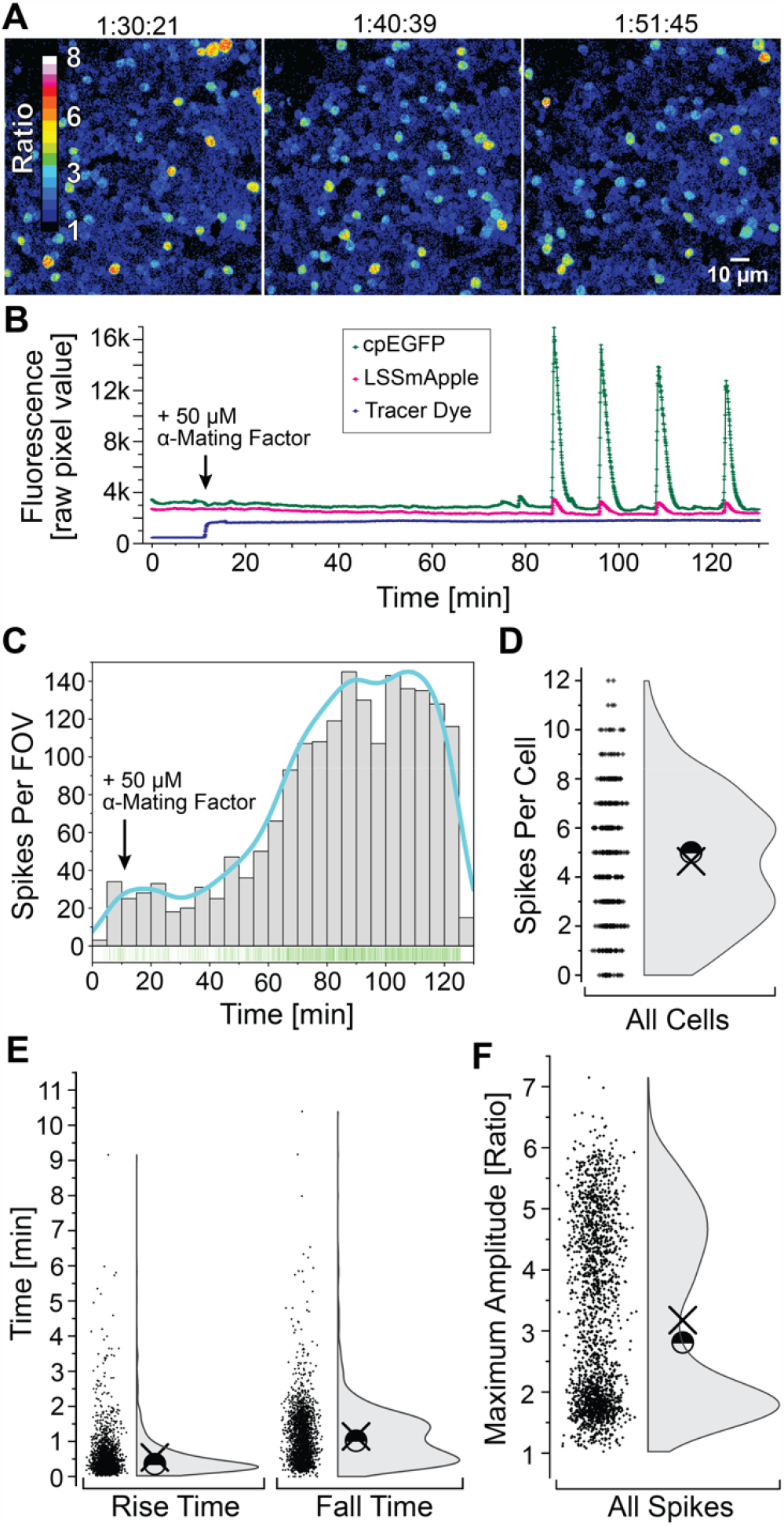
Calcium spiking in S. *cerevisiae* GA-MatryoshCaMP6s lines. **(A)** Average z-stack projections of confocal images of yeast expressing GA-MatryoshCaMP6s after α-MF treatment. Ratio (cpEGFP/LSSmApple) values are shown as the ratio of green to red fluorescence intensity (FI) by 16-color lookup table. Five independent biological replicates were analyzed with similar results. Time stamps are in hour:minute:second format. **(B)** Sensor response from a single yeast cell. Graph displays the mean raw pixel FI values for green and red channels for the GA-MatryoshCaMP6s, as well tracer dye in blue. Arrowhead indicates the timepoint in minutes (min) of the treatment (50 μM α-MF). **(C)** Histogram depicts number of calcium spikes in field of view (FOV) over time. Cyan line shows kernel density estimate smoothing function. Dark green lines indicate spike count (n=1898). **(D)** Number of spikes per cell (all cells n=409). Half circle marks the median (5) and chi symbol the mean (4.64±0.13). **(E)** Rise and fall time of all spikes (n=1898). Median (rise time 0.35 min, fall time 1.02 min) Mean (rise time 0.54±0.04 min, fall time 1.14±0.02 min). **(F)** Maximum changes in ratio (cpEGFP/LSSmApple) of all spikes (n=1898). Median (2.81) and mean ± standard error (3.18±0.03).

## Discussion

The newly engineered LSSmApple FP has spectral properties that facilitate multiplexing with green FPs, such as cpEGFP, which has been used extensively in construction of fluorescent sensors. We created a nested fusion of cpEGFP and LSSmApple and used the cassette to convert GCaMP6s into a monochromatically excitable ratiometric calcium indicator. We expressed the GA-MatryoshCaMP6s sensor in budding yeast to visualize responses to α-MF. Cytosolic calcium elevations have long been implicated in α-MF responses^22,24^. Results from population-scale assays using radiolabelled calcium (^45^Ca^2+^) or the chemiluminescent calcium indicator aequorin prompted earlier researchers to conclude there is only a single calcium elevation during the yeast mating response^25^. We clearly observed asynchronous, repetitive calcium spiking, consistent with a more recent study that concluded that α-MF treatment strongly increased frequency of calcium spiking^21^. When averaged across a large population, asynchronous spiking would appear to be single, longer-duration event, which highlights the importance of single-cell analysis. Taking advantage of LSSmApple as a stable reference channel, we developed an analysis pipeline to segment and track individual cells to facilitate quantitative analysis. With our approach, hundreds of cells can be individually analyzed, and the same materials and methods can be readily adapted to other sensors or other treatments reported or posited to elicit calcium signals in yeast. Indeed, because of the simplicity of cloning and imaging GA-Matryoshka sensors, we expect that others will adopt the technology to facilitate ratiometric measurements, segmentation, and tracking. The AI-assisted data analysis pipeline that we developed for segmentation and tracking of yeast cells demonstrates the added value of incorporating a stable reference FP in intensiometric sensors and could see a host of applications in yeast cell biology and fluorescent sensor characterization.

## Conclusion

LSSmApple is a new LSS FP that is advantageous for use in combination with widely used green FPs, including cpEGFP. By nested fusion of LSSmApple in cpEGFP, we engineered a second-generation “Green-Apple Matryoshka” cassette that enables rapid conversion of intensiometric cpEGFP sensors into dual-FP sensors with a stable reference channel that can be used for ratiometric measurements, as well as segmentation and tracking of regions of interest. For faithful ratiometric readouts, researchers are advised to use image acquisition parameters that minimize photobleaching, as achieved here, because divergent photobleaching rates would distort ratiometric values. While a direct comparison with GO-Matryoshka was not conducted, we anticipate that the ease of co-excitation, compatibility with standard RFP filters, and comparable maturation times between the two fluorophores of the GA-Matryoshka sensor will propel the widespread. We used the GA-Matryoshka cassette to create GA-MatryoshCaMP6s and performed calcium imaging in yeast. The software pipeline we created enables segmentation and tracking of hundreds of cells using the LSSmApple channel. While employed for mere proof of concept here, we anticipate that our analysis pipeline, GA-MatryoshCaMP6s, and future GA-Matryoshka sensors will be powerful and accessible tools for yeast biologists. More broadly, GA-Matryoshka technology should provide a widely useful platform for fluorescent sensor engineering.

## Material and methods

### LSSmApple engineering and characterization

All synthetic DNA oligonucleotides for cloning and library construction were purchased from Integrated DNA Technologies (IDT). *Taq* DNA polymerase (New England Biolabs) was used for error-prone PCR (EP-PCR). All site-directed mutagenesis were performed using the Quikchange lightning mutagenesis kit (Agilent) and primers designed according to the manufacturer’s guidelines. PCR products and products of restriction digests were purified using gel extraction kit (Thermo) according to the manufacturer’s protocols. Restriction enzymes and ligases were purchased from New England Biolabs or Thermo Scientific. The cDNA sequences were confirmed by dye terminator cycle sequencing using the BigDye Terminator v3.1 Cycle Sequencing Kit (Applied Biosystems). Sequencing reactions were analyzed at the University of Alberta Molecular Biology Service Unit.

Engineering of LSSmApple was carried out by site-directed mutagenesis and iterative rounds of EP-PCR, starting from the mApple gene as the initial template. EP-PCR products were digested with XhoI and *HindIII* and ligated into pBAD/His B vector digested with the same two enzymes. The ligation product was used to transform electrocompetent *E. coli* strain DH10B (Invitrogen), which were then plated on agar plates containing LB medium supplemented with 0.4 mg/mL ampicillin and 0.02 % w/v L-arabinose. Single colonies were picked and inoculated into 4 mL of LB medium with 0.1 mg/mL ampicillin and 0.02 % w/v L-arabinose and then cultured overnight. Protein was extracted using B-PER bacterial extraction reagent (Thermo Fisher Scientific). Libraries were screened for variants that exhibited brighter LSS red fluorescence in colonies of *E. coli* using a custom-built fluorescent imaging system equipped with filter sets (Chroma) for 470/40 nm excitation with 630/60 nm emission^26^. Colonies exhibiting the highest intensity of LSS fluorescence were picked and cultured for further spectral confirmation.

To purify FP variants for characterization, electrocompetent *E. coli* strain DH10B (Invitrogen) was transformed with the plasmid of interest using a Micropulser electroporator (Bio-Rad). Bacteria were cultured overnight on agar plates containing LB and ampicillin. Single colonies were picked and grown overnight in 4 mL LB supplemented with ampicillin at 37 °C. For each colony, the 4 mL culture was then used to inoculate 250 mL of LB medium with ampicillin and grown to an optical density of OD_600_ = 0.6. Protein expression was induced with the addition of 0.02 % L-arabinose and the culture was grown overnight at 37 °C, 220 rpm. Bacteria were pelleted at 10,000 rpm, 4 °C for 10 min, lysed using a cell disruptor (Constant Systems) and the suspension was clarified at 14,000 rpm for 30 min. The protein was purified from the supernatant by Ni-NTA affinity chromatography (ABT) according to the manufacturer’s instructions. The buffer of the purified protein was exchanged with 10 mM Tris-Cl, 150 mM NaCl, and pH 7.3 with Amicon ultra centrifugal filters (MWCO 10,000) for a final protein concentration of approximately 10 μM.

Molar extinction coefficients (EC) were measured by the alkali denaturation method^27^. Briefly, the protein was diluted into Tris buffer or 1 M NaOH and the absorbance spectra were recorded under both conditions. The EC was calculated assuming the denatured RFP chromophore has an EC of 44,000 M^-1^cm^-1^ at 452 nm. Fluorescence quantum yields (QY) were determined using LSSmKate2 as a standard^27^. Fluorescence intensity as a function of pH was determined by dispensing 2 μL of the protein solution into 50 μL of the desired pH buffer in triplicate into a 384-well clear-bottomed plate (Nunc) and measured in a Safire2 plate reader (Tecan). pH Buffer solutions from pH = 3 to pH = 11 were prepared according to the Carmody buffer system^28^. Plasmid containing LSSmApple is available from AddGene (Plasmid #191259). LSSmApple spectral data have been uploaded to FPbase and are accessible at https://www.fpbase.org/protein/lssmapple/.

### Cloning of GA-MatryoshCaMP6s constructs

GA-MatryoshCaMP6s was constructed by replacement of the coding sequence of LSSmOrange in GO-MatryoshCaMP6s with the LSSmApple coding sequence by In-Fusion cloning (Takara) following manufacturer recommendations. The resulting constructs were sub-cloned into pRSET B for protein expression in *E. coli* BL21 (DE3), adding an N-terminal His-6-tag to the constructs for purification. For yeast transformation, GA-MatryoshCaMP6s was amplified from pRSET B His-GA-MatryoshCaMP6s and subcloned into pDONR /Zeo via In-Fusion cloning (Takara Bio, GA-MatryoshCaMP6s forward primer: ccaactttgtacaaaaaagcaggcttaatggtcgactcatcacgtcg, GA-MatryoshCaMP6s reverse primer: caagaaagctggttttatcacttcgctgtcatcatttgtac). Subsequently, GA-MatryoshCaMP6s was transferred into the Destination vector pAG416 GPD via LR cloning. (pAG416 GPD: https://dx.doi.org/10.1002%2Fyea.1502)^29^. Sequences were verified by whole-plasmid sequencing (Plasmidsaurus). Plasmid containing GA-MatryoshCaMP6s will be deposited in AddGene and is available upon request.

### GA-MatryoshCaMP6s expression and purification

*E. coli* BL21 (DE3) (New England Biolabs C2527H) was transformed with pRSET B-GA-MatryoshCaMP6s. As starter culture single colonies were inoculated 14-18 h in 2-3 mL Luria broth (LB) with 50 μg/mL carbenicillin disodium (Duchefa Biochemie; CAS 4800-94-6) in glass tubes, at 37 °C, 200 rpm. 2mL of the starter cultures were transferred into a final volume of 100 mL LB with 50 μg/mL carbenicillin disodium, 0.2 % D-Lactose (Sigma Aldrich; CAS 64044-51-5) and 0.05 % D-Glucose monohydrate (Thermo Scientific Chemicals; CAS 14431-43-7) in 500 mL Erlenmeyer flasks without baffles. After 2 h incubation on 37 °C at 200 rpm, cultures were transferred to 20 °C at 200 rpm and incubated for 48 h. Bacterial cultures were cooled on ice and centrifuged for 10 min at 3790 g and 4 °C. The pellet was stored at -20 °C at least 14 h, after unfreezing on ice the pellet was resuspended in 1.5 mL 20 mM MOPS (Roth; CAS 1132-61-2) pH 7.0. To prevent sensor degradation 1 cOmpleteTM ULTRA Tablet Mini protease inhibitor cocktail (Roche) was used per 100 mL of all purification buffers. Cells were opened via sonication (QSonica Q700 Sonicator) with Amplitude 50, 45 s process time, 3 s pulse on and 8 s pulse off time. The sonicated sample was centrifuged 10 min at 15871 g, 4 °C to remove cellular debris. Purification of His-tagged GA-MatryoshCaMP6s was performed with Ni-NTA agarose beads (Qiagen, cat. No: 30210). Before applying the lysate, 1.5 mL Ni-NTA agarose beads were loaded on a column and washed with 5-10 ml 20 mM MOPS. After loading of the lysate another wash step with 5-10 ml 20 mM MOPS was performed. His-tagged GA-MatryoshCaMP6s were eluted in 1.5 mL 250 mM imidazole (Sigma-Aldrich; CAS 1467-16-9) 20 mM MOPS pH 7.0. Protein concentration was measured in NanoDrop™ One (Thermo Scientific). Purified sensors were incubated at 4 °C at least 14 h to ensure maturation of the fluorescent protein.

### Fluorimetric analysis of GA-MatryoshCaMP6s

For the calcium titrations, Commercial calcium Calibration Buffer Kit #1 (Invitrogen Life Technology, Paisley, United Kingdom) was used. The stock solutions of zero-free calcium buffer (10 mM K_2_EGTA, 100 mM KCl, 30 mM MOPS, pH 7.2) and 39 μM calcium buffer (10 mM Ca-EGTA, 100 mM KCl, 30 mM MOPS, pH 7.2) were mixed according to the manufacturer, yielding 11 different free calcium concentrations. Purified sensors were set to a concentration of 5 mg/mL and then diluted 1:10 in zero-free calcium buffer, to buffer residual calcium in the elution buffer (250 mM imidazole 20 mM MOPS pH 7.0). Per well, 10 μL of sensor were mixed with 190 μL of calcium buffer. Final calcium concentrations in the well were calculated with Ca-EGTA Calculatorv1.2^30^ (https://somapp.ucdmc.ucdavis.edu/pharmacology/bers/maxchelator/CaEGTA-NIST.htm) (free calcium at 27 °C, pH 7.2, 0.1 N: 2280 nM, 837.7 nM, 443.7 nM, 276.4 nM, 184 nM, 125.3 nM, 84.8 nM, 55 nM, 32,4 nM, 14,5 nM, 0 nM zero-free calcium). Ligand titrations were performed by using a microplate reader TECAN™ Spark^®^, for cpEGFP point measurement, λ_ex_ 480/10 nm; λ_em_ 515/10 nm and for LSSmApple point measurement, λ_ex_ 460/10 nm; λ_em_ 600/10 nm with Dichroic 510 mirror. Steady-state fluorescence spectra were measured using λ_ex_ 453/10 nm and λ_em_ 499-700/10 nm with Δλ 2 nm step size with a 50% mirror. All measurements were recorded in top reading mode with 80-130 gain and temperature controlled at 27±0.5 °C. Proteins were used at a final concentration of 25 μg/mL. Spectra were background-subtracted using a zero-free calcium buffer control. Analysis was performed in 96-well, transparent, flat-bottom, half-area plates (Greiner Bio-One, Germany). The fluorescence emission intensity ratios before adding ligand (R_0_) and with increasing ligand concentration (R) were calculated by dividing the fluorescence intensity of cpEGFP (λ_ex_ 480 nm, λ_em_ 515 nm) by the intensity of LSSmApple (λ_ex_ 460 nm, λ_em_ 600 nm). Data was normalized accordingly:

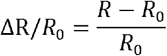

Analysis and visualization of the data was performed using Excel, Origin Lab 2020 and Illustrator 2020. Sigmoid fits were done using MyCurveFit (https://mycurvefit.com). Data were entered in mean and standard error of ΔR/R_0_ of n=3 biological replicates (n=3 technical replicates per biological replicates) out of two independent titrations. We proceeded in the same way for ΔF/F_0_. K_d_ was determined as ligand concentration at 50 % dynamic range (ΔR/R_0_ or ΔF/F_0_) in sigmoid function.

### Homology modeling of proteins

Homology models for LSSmApple, the GA-Matryoshka cassette and the GA-MatryoshCaMP6s were generated with AlphaFold2 (ColabFold v1.5.2: AlphaFold2 using MMseqs2)^31–34^. The models with the best ranked pLDDT were chosen for display. 3D-visualization of protein structures was performed using the molecular visualization system PyMOL 2.5.4 (Schrödinger). For better visualization of the chromophore of LSSmApple in the right panel of Figure 1 A, an N-terminal α-helix (amino acid sequence: VSKGEENNMAI) and a C-terminal α-helix (amino acid sequence: MDELYK) were hidden with PyMOL. The model’s confidences (pLDDT values from AlphaFold2) were visualized via b-factor spectrum of PyMOL in Figure S1.

### Yeast growth, genetic transformation, and trapped-cell microfluidics

Yeast strain K601/ W303-1A MATa *(ade2-1 ura3-1 his3-11 trp1-1 leu2-3 leu2-112 can1-100)* was a gift from Cristina Bonza^35^. Preparation of competent yeast cells and polyethylene glycol (PEG) 3350-mediated transformation were conducted as described previously^36^. K601 was transformed with pAG416-GPD encoding GA-MatryoshCaMP6s. pAG416GPD-ccdB was a gift from Susan Lindquist (Addgene plasmid #14148; http://n2t.net/addgene:14148 ; RRID:Addgene_14148) A minimum of 3 glycerol stocks was prepared from independently transformed colonies in YPAD 25% glycerol. K601 GA-MatryoshCaMP6s strains were grown on SC-Ura medium (For 1 L: CSM-Ura 0.77 g YNB w/ ammonium 6.7 g, glucose 20 g., agar 20 g (with adenine addition, total concentration of adenine 150 μM)). For trapped cells microfluidics, glycerol stocks were streaked out on SC-Ura plates and incubated at 29.5 °C for 2 days. Colonies were resuspended in liquid SC-Ura at OD_600_ = 0.5-1 and loaded into CellASIC ONIX plate for haploid yeast cells (Cat#Y04C-02-5PK). CellASIC ONIX plates were accordingly loaded with SC-Ura as mock and 50 μM α-MF in SC-Ura (Bachem AG (Cat# 4047325)) for treatment. Plates were sealed with CellASIC ONIX 2 temperature-controlled manifold (Cat# CAX2-MXT20) and connected to the CellASIC ONIX 2 live-cell imaging platform (Cat# CAX2-S0000). Trapped yeast cells were incubated overnight with a constant perfusion of SC-Ura driven by 17 kPa of pressure at 28 °C. Before starting experiments, cells were acclimated to a perfusion of SC-Ura driven by 20 kPa. Experiments were conducted at 28 °C with a constant perfusion driven by 20 kPa pressure and included a 10 min perfusion with SC-Ura (mock) followed by perfusion with 50 μM α-MF in SC-Ura medium containing 1 ^μ^g/mL ATTO 643 as tracer dye (ATTO-TEC GmbH).

### Fluorescence microscopy

Quantitative calcium imaging of *S. cerevisiae* GA-MatryoshCaMP6s was performed using an Olympus IXplore SpinSR spinning disk confocal microscope. Samples were excited with a 488 nm laser (Obis, 100 mW, used at 1-2% power) for GA-MatryoshCaMP6s and with a 640 nm laser (Obis, 100 mW, used at 1-2% laser power) for ATTO 643. The microscope was equipped with a Yokogawa confocal scanning unit (CSU)-W1 SoRa micro-lensed pinhole disk and an Olympus UAPON 100× OTIRF oil-immersion objective (NA 1.49). Emission intensities were collected in rapid sequential stacks using a 525/50 nm emission filter for cpEGFP, 617/73 nm emission filter for LSSmApple, and 685/40 nm emission filter for the ATTO 643 tracer dye. For detection, an Andor iXon Ultra 888 electron-multiplying charge coupled device (EMCCD) was used. Z-stacks were performed by using a Mad City Labs Z-axis piezo nanopositioner with 300 nm travel range (Cat# OLY-S1023-Nano-ZL300-OSSU). Z-drift compensation (Olympus) was used to maintain focal plane during acquisition. Full Z-stack acquisitions for each channel were performed with a frame rate of 3 s. Exposure time for each channel was 20 ms, with 4x4 pixel binning.

### Image analysis

During the first processing steps in Fiji [38] (V 1.53s), original channels remained unprocessed, except that the image size in pixel was doubled for annotation and quality control reasons. All channels were treated the same. In addition, a fourth channel was generated as a Fiji “walking average” taking 20 frames into the LSSmApple-channel into account and was subsequently processed using the Fiji function of “Enhance local contrast” for two times, to create and highlight an intracellular signal of similar intensity for nearly all of the cells that would be used for segmentation in the next step. Second, the data were used in a CellProfiler [39] (V 4.2.1) workflow. Segmentation was performed by utilizing the “RunCellpose” plugin” (Cellpose V.0.6.5) on the previously created fourth segmentation channel of the dataset and resulted in segmented yeast for the whole timeseries. The pretrained Cellpose nucleus network was used with an expected object diameter of 16 was set and a flow threshold of standard 0.4 and a cell probability threshold of 0 was used. To track and link generated segmented objects in time, the track objects feature of CellProfiler was used. The “distance” algorithm was chosen as tracking method with a minimum considered distance of 1 pixel. The results tables of cellular features and intensities over time were exported for later use in KNIME. Annotated graphical outputs were additionally generated to visually check and ensure quality of the segmentation and tracking. Third, time tracked data were sorted and classified in KNIME [40] (V 4.5.2) based on the previously extracted features. Again, as a quality control, graphical outputs were generated that allowed backtracking of specific tracks to the original timeseries / segmentation. Furthermore, cells that could not be continuously tracked for at least 90% of the frames in the timeseries were rejected. For each track the intensities of the cpEGFP and the LSSmApple channel were isolated as mean intensity 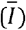 and the ratio (R_int_) at every timepoint was calculated for every cell by dividing intensities of cpEGFP by LSSmApple. A baseline ratio (R_med_) of each track was calculated as the median of all ratios of a cell over time and the maximum peak ratio (R_max_) observed for one cell was identified. The difference (Δ_*R*_) between R_med_ and R_max_ was used to classify the response strength of each cell into 3 different classes. 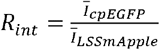 and 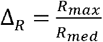 with

R_int_ = ratio between intensities of cpEGFP readout and LSSmApple reference

I = mean intensity of the corresponding channel per cell in grey values

R_med_ = median of all R_int_ per cell found over time (baseline ratio)

R_max_ = maximum value of all R_int_ found per cell over time (peak maximum ratio)

ΔR = difference between median baseline and peak maximum (peak ratio amplitude)

Tracks below a ΔR of 0.6 were considered as cells with no significant response, tracks with a ΔR between 0.6 and 2 as cells with a weak response and cells with ΔR of more than 2 as strong responding cells. Finally, data of these classified cells as ratio over time were exported and the percentage contribution of each of the classes to the total cell count was calculated^37–40^. The resulting data were exported for graph preparation in Origin Pro 2020. Half-violin plots and histogram smoothing function show kernel density estimates. Workflow files are accessible at: https://github.com/SHaensch/2022_YIRAS_YeastIntensity RatioAndSegmentation^38–40^.

## ASSOCIATED CONTENT

### Supporting Information

Confidence maps of AlphaFold2 models; GA-MatryoshCaMP6s response to α-MF in yeast,.

## Author Contributions

Conception: JOE, MS, YS, WBF, REC, TJK. Data collection: JOE, MS, YS, YI, ES, TJK. Data analysis and interpretation: all authors. Supervision: MS, YI, WBF, MMW, REC, TJK. Writing of the initial draft and figure preparation: JOE, MS, YS, MMW, TJK. Manuscript editing and contributions: all authors. Funding: WBF, REC.

## ACKNOWLEDGMENTS

The authors thank Susanne Paradies for technical assistance and Marcel Dickmanns for helping with AlphaFold2. The authors would also like to thank Prof. Dr. Guido Grossmann for helpful comments on project design and presentation.

## ABBREVIATIONS

Ca^2+^: calcium
GA: green-apple
GO: green-orange
EGFP: enhanced green fluorescent protein
mApple: monomeric Apple
LSS: large Stokes shift
QY: quantum yield
FRET: Förster resonance energy transfer
SNR: signal-to-noise ratio
cp: circularly permutated
ESPT: excited state proton transfer
E: glutamic acid
D: aspartic acid
FI: fluorescent intensity
ROI: region of interest
FOV: field of view
EP-PCR: error-prone PCR
EC: extinction coefficient
SDM: site-directed mutagenesis
SC-Ura: synthetic complete media without Uracil
α-MF: α-mating factor
nm: nanometer
YPAD: yeast extract peptone adenine dextrose medium
PEG: polyethylene glycol
s. e. m.: standard error of mean.

## References

1. Chalfie, M., Tu, Y., Euskirchen, G., Ward, W. W. & Prasher, D. C. Green Fluorescent Protein as a Marker for Gene Expression. Science (1979) 263, 802–805 (1994).

2. Shimoura, O., Johnson, F. H. & Saiga, Y. Extraction, purification and properties of Aequorin, a bioluminescent Protein from the Luminous Hydromedusan, Aequorea. J Cell Comp Physiol 59, 223–239 (1962).

3. Prasher, D. C., Eckenrode, V. K., Ward, W. W., Prendergast, F. G. & Cormier, M. J. Primary structure of the Aequorea victoria green-fluorescent protein. Gene 111, 229–233 (1992).

4. Rodriguez, E. A. et al. The Growing and Glowing Toolbox of Fluorescent and Photoactive Proteins. Trends Biochem Sci 42, 111–129 (2017).

5. Lambert, T. J. FPbase: a community-editable fluorescent protein database. Nat Methods 16 277–278 (2019).

6. Stokes, G. G. On the change of refrangibility of light. Philos Trans R Soc Lond 142, 463–562 (1852).

7. Chu, J. et al. A bright cyan-excitable orange fluorescent protein facilitates dual-emission microscopy and enhances bioluminescence imaging in vivo. Nat Biotechnol 34, 760–767 (2016).

8. Shcherbakova, D. M., Hink, M. A., Joosen, L., Gadella, T. W. J. & Verkhusha, V. V. An orange fluorescent protein with a large stokes shift for single-excitation multicolor FCCS and FRET imaging. J Am Chem Soc 134, 7913–7923 (2012).

9. Shen, Y., Chen, Y., Wu, J., Shaner, N. C. & Campbell, R. E. Engineering of mCherry variants with long Stokes shift, red-shifted fluorescence, and low cytotoxicity. PLoS One 12, (2017).

10. Frommer, W. B., Davidson, M. W. & Campbell, R. E. Genetically encoded biosensors based on engineered fluorescent proteins. Chem Soc Rev 38, 2833–2841 (2009).

11. Deuschle, K. et al. Rapid metabolism of glucose detected with FRET glucose nanosensors in epidermal cells and intact roots of Arabidopsis RNA-silencing mutants. Plant Cell 18, 2314–2325 (2006).

12. Kaper, T., Lager, I., Looger, L. L., Chermak, D. & Frommer, W. B. Fluorescence resonance energy transfer sensors for quantitative monitoring of pentose and disaccharide accumulation in bacteria. Biotechnol Biofuels 1, (2008).

13. Sadoine, M. et al. Designs, applications, and limitations of genetically encoded fluorescent sensors to explore plant biology. Plant Physiol 187, 485–503 (2021)

14. Baird, G. S., Zacharias, D. A. & Tsien, R. Y. Circular permutation and receptor insertion within green fluorescent proteins. Proceedings of the National Academy of Sciences 96, 11241–11246 (1999).

15. Nagai, T., Sawano, A., Park, E. S. & Miyawaki, A. Circularly permuted green fluorescent proteins engineered to sense Ca^2+.^Proceedings of the National Academy of Sciences 98, 3197–3202 (2001).

16. Walia, A., Waadt, R. & Jones, A. M. Genetically Encoded Biosensors in Plants: Pathways to Discovery. Annu Rev Plant Biol 69, 497–524 (2018).

17. Ast, C. et al. Ratiometric Matryoshka biosensors from a nested proteins. Nat Commun 8, 431 (2017)

18. Piatkevich, K. D. et al. Monomeric red fluorescent proteins with a large Stokes shift. Proceedings of the National Academy of Sciences 107, 5369–5374 (2010).

19. Pletnev, S. et al. Orange fluorescent proteins: Structural studies of LSSmOrange, PSmOrange and PSmOrange2. PLoS One 9, (2014).

20. Chen, T. W. et al. Ultrasensitive fluorescent proteins for imaging neuronal activity. Nature 499, 295–300 (2013).

21. Carbó, N., Tarkowski, N., Ipiña, E. P., Dawson, S. P. & Aguilar, P. S. Sexual pheromone modulates the frequency of cytosolic Ca^2+^bursts in Saccharomyces cerevisiae. Mol Biol Cell 28, 501–510 (2017).

22. Hohmann, S. Osmotic Stress Signaling and Osmoadaptation in Yeasts. Microbiology and Molecular Biology Reviews 66, 300–372 (2002).

23. Bermejo, C., Haerizadeh, F., Takanaga, H., Chermak, D. & Frommer, W. B. Optical sensors for measuring dynamic changes of cytosolic metabolite levels in yeast. Nat Protoc 6, 1806–1817 (2011).

24. Iida, H., Yagawa, Y. & Anraku, Y. Essential role for induced Ca^2+^ influx followed by [Ca^2+^](i) rise in maintaining viability of yeast cells late in the mating pheromone response pathway. A study of [Ca^2+^](i) in single Saccharomyces cerevisiae cells with imaging of fura-2. Journal of Biological Chemistry 265, 13391–13399 (1990).

25. Nakajima-Shimada, J., Sakaguchi, S., Tsuji, F. I., Anraku, Y. & Iida, H. Ca^2+^signal is generated only once in the mating pheromone response pathway in Saccharomyces cerevisiae. Cell Struct Funct 25, 125–131 (2000).

26. Ai, H. W., Baird, M. A., Shen, Y., Davidson, M. W. & Campbell, R. E. Engineering and characterizing monomeric fluorescent proteins for live-cell imaging applications. Nat Protoc 9, 910–928 (2014).

27. Cranfill, P. J. et al. Quantitative assessment of fluorescent proteins. Nat Methods 13, 557–562 (2016).

28. Carmody, W. R. Easily prepared wide range buffer series. J Chem Educ 38, 559 (1961).

29. Alberti, S., Gitler, A. D. & Lindquist, S. A suite of Gateway® cloning vectors for high-throughput genetic analysis in Saccharomyces cerevisiae. Yeast 24, 913–919 (2007).

30. Bers, D. M., Patton, C. W. & Nuccitelli, R. A practical guide to the preparation of Ca^2+^buffers. Methods in Cell Biology vol. 99 (2010).

31. Mitchell, A. L. et al. MGnify: the microbiome analysis resource in 2020. Nucleic Acids Res. 48, D570–D578 (2019)

32. Mirdita, M. et al. Uniclust databases of clustered and deeply annotated protein sequences and alignments. Nucleic Acids Res. 45, D170–D176 (2017).

33. Mirdita, M., Steinegger, M. & Söding, J. MMseqs2 desktop and local web server app for fast, interactive sequence searches. Bioinformatics 35, 2856–2858 (2019).

34. Mirdita, M. et al. ColabFold: Making Protein folding accessible to all. Nat Methods 19, 679–682 (2022).

35. Bonza, M. C., Luoni, L. & De Michelis, M. I. Functional expression in yeast of an N-deleted form of At-ACA8, a plasma membrane Ca^2+^-ATPase of Arabidopsis thaliana, and characterization of a hyperactive mutant. Planta 218, 814–823 (2004).

36. Gietz, R. D. & Schiestl, R. H. High-efficiency yeast transformation using the LiAc/SS carrier DNA/PEG method. Nat Protoc 2, 31–34 (2007).

37. Schindelin, J. et al. Fiji: An open-source platform for biological-image analysis. Nat Methods 9, 676–682 (2012).

38. Stirling, D. R. et al. CellProfiler 4: improvements in speed, utility and usability. BMC Bioinformatics 22, 433 (2021).

39. Stringer, C., Wang, T., Michaelos, M. & Pachitariu, M. Cellpose: a generalist algorithm for cellular segmentation. Nat Methods 18, 100–106 (2021).

40. Berthold, M. R. et al. KNIME - the Konstanz Information Miner: Version 2.0 and Beyond. SIGKDD Explor. Newsl. 11, 26–31 (2009).

