## Supplementary Information for "A monochromatically excitable green-red dual-fluorophore fusion incorporating a new large Stokes shift fluorescent protein"

**SUPPORTING INFORMATION**

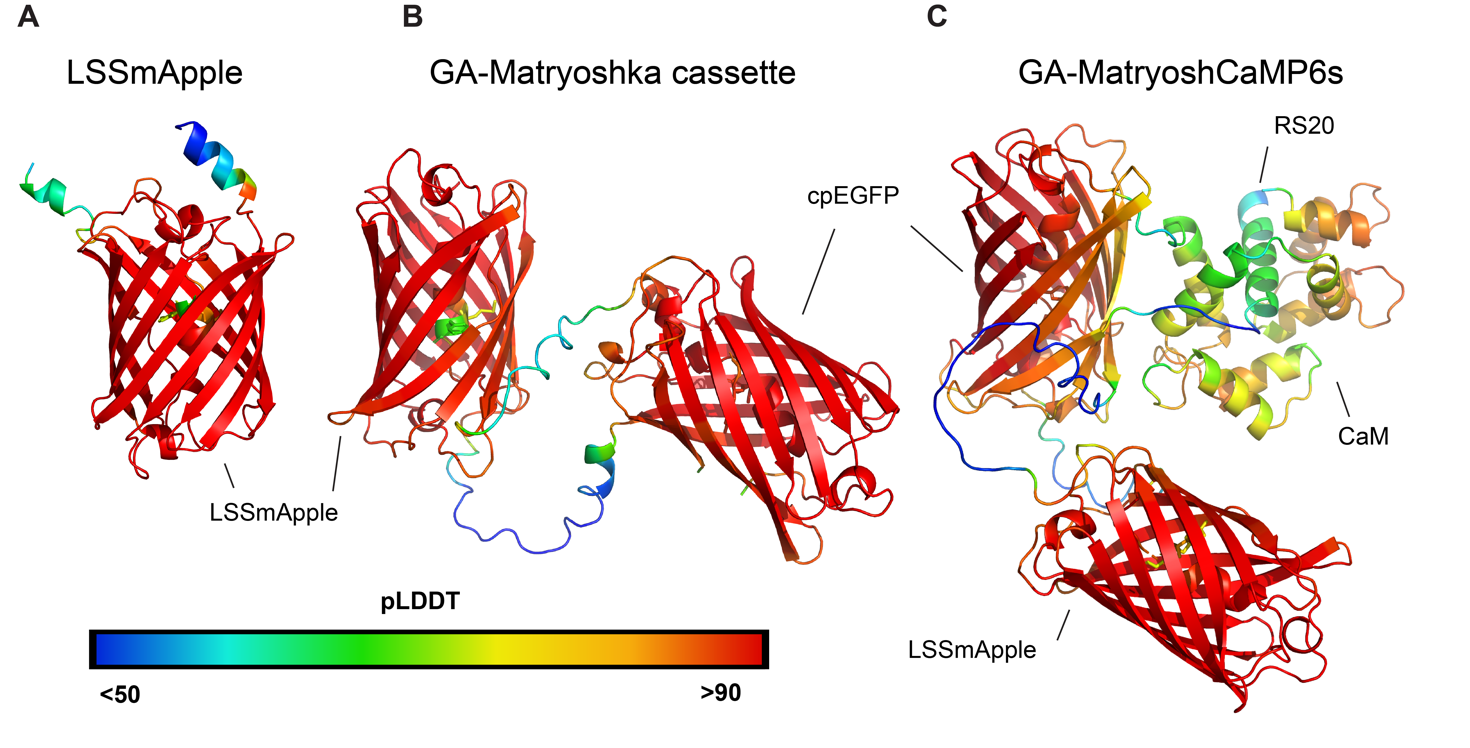

Figure S1. Confidence of AlphaFold2 models. AlphaFold2 models of GA-Matryoshka cassette and GA-MatryoshCaMP6s colored after confidence of the prediction. High predicted local distance difference test (pLDDT) values (yellow to red) correspond to high confidence and low pLDDT values (blue to green) to lower confidence.

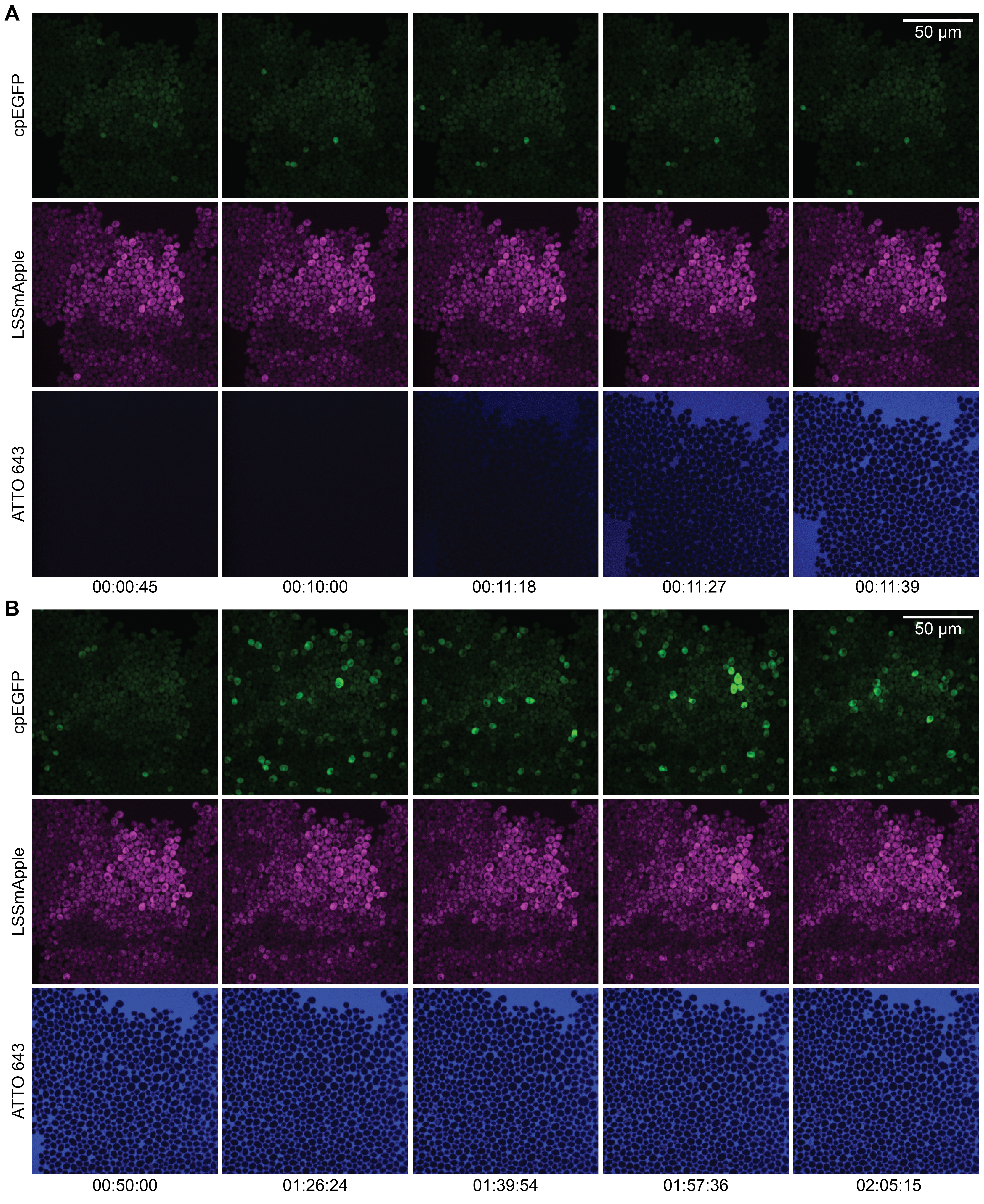

Figure S2. GA-MatryoshCaMP6s response induced by treatment of yeast with α-MF. **(A)** Panel shows yeast cells before and after perfusion of 50 µM α-MF. **(B)** Asynchronous calcium spiking in response to prolonged treatment with α-MF. Images of cpEGFP fluorescence are shown in the top row of images. LSSmApple fluorescence is shown in the middle row. The bottom row shows a far-red tracer dye (ATTO 643) labeling treatment solution. Time stamps are in hour:minute:second format.

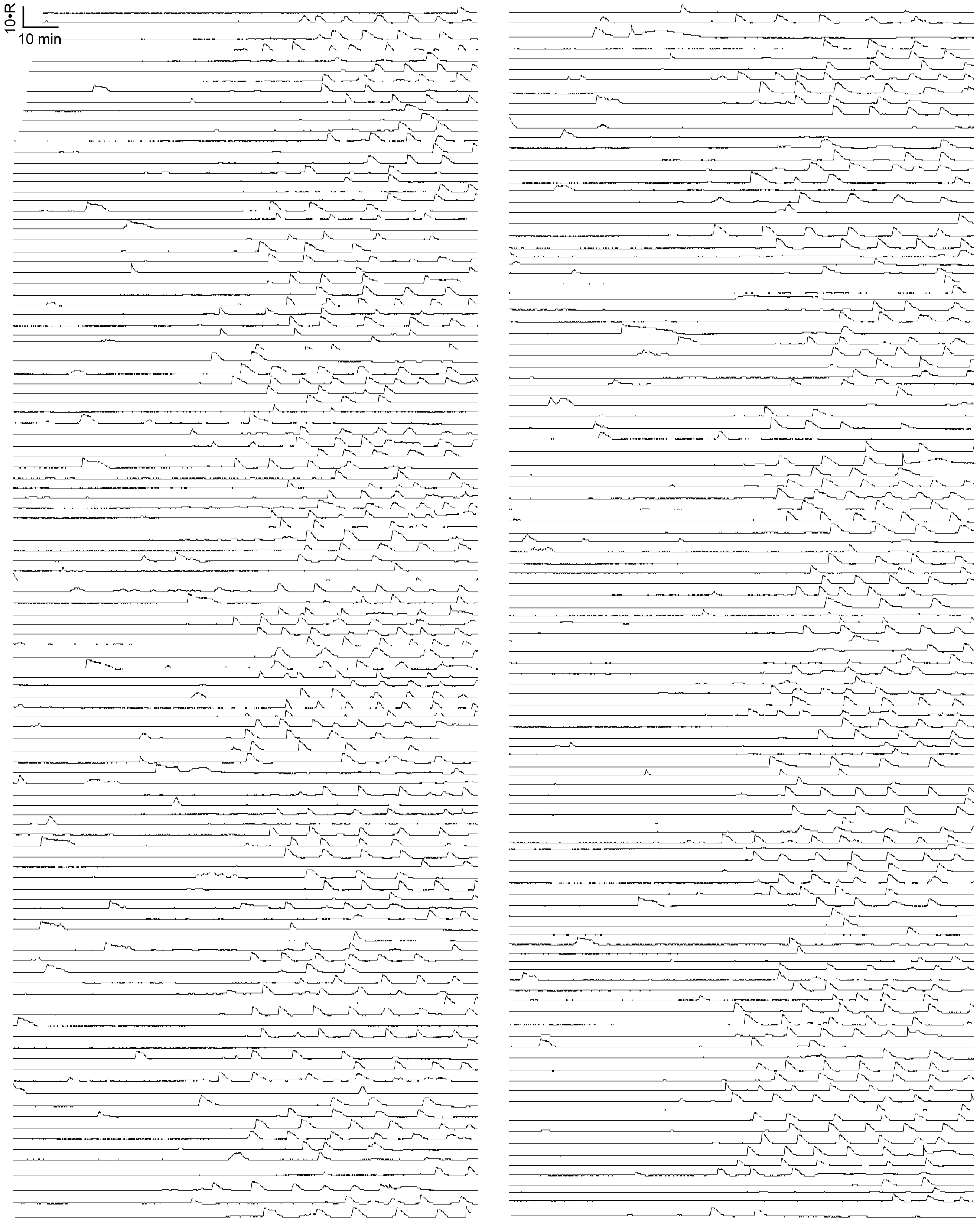

Figure S3. Traces of single-cell regions of interest (ROIs) from a single α-MF perfusion experiment. Scale bars indicate 10 minutes (min) and a tenfold change in cpEGFP / LSSmApple fluorescence intensity ratio (10•R).
